# *De novo* protein identification in mammalian sperm using high-resolution *in situ* cryo-electron tomography

**DOI:** 10.1101/2022.09.28.510016

**Authors:** Zhen Chen, Momoko Shiozaki, Kelsey M. Haas, Shumei Zhao, Caiying Guo, Benjamin J. Polacco, Zhiheng Yu, Nevan J. Krogan, Robyn M. Kaake, Ronald D. Vale, David A. Agard

## Abstract

Understanding molecular mechanisms of cellular pathways requires knowledge of the identities of participating proteins, their cellular localization and their 3D structures. Contemporary workflows typically require multiple techniques to identify target proteins, track their localization using fluorescence microscopy, followed by *in vitro* structure determination. To identify mammal-specific sperm proteins and understand their functions, we developed a visual proteomics workflow to directly address these challenges. Our *in situ* cryo-electron tomography and subtomogram averaging provided 6.0 Å resolution reconstructions of axonemal microtubules and their associated proteins. The well-resolved secondary and tertiary structures allowed us to computationally match, in an unbiased manner, novel densities in our 3D reconstruction maps with 21,615 AlphaFold2-predicted protein models of the mouse proteome. We identified Tektin 5, CCDC105 and SPACA9 as novel microtubule inner proteins that form an extensive network crosslinking the lumen of microtubule and existing proteins. Additional biochemical and mass spectrometry analyses helped validate potential candidates. The novel axonemal sperm structures identified by this approach form an extensive interaction network within the lumen of microtubules, suggesting they have a role in the mechanical and elastic properties of the microtubule filaments required for the vigorous beating motions of flagella.

## Introduction

The beating motion of sperm flagella is essential for natural fertilization^1^. All cilia and flagella share a conserved filamentous structure, the axoneme that contains nine doublet microtubules (doublets) surrounding two singlet microtubules^2,3^. However, mammalian sperm microtubules have to withstand larger bending torques compared to other motile cilia^4,5^. Furthermore, sperm from different species can differ dramatically in their morphologies and swimming behaviors. Despite their importance in fertility and speciation, we have limited understanding of the unique adaptations of sperm axonemes at the molecular level. In this study, we combined high-resolution *in situ* cryoET and AlphaFold2 modeling in a visual proteomics workflow to directly identify, without labeling, novel microtubule-associated proteins in mouse and human sperm and their structures and interactions in cells.

### *In situ* structures of sperm doublets at subnanometer resolutions

To suppress the conformational dynamics of the sperm axonemes, freshly extracted mouse sperm were treated with the dynein ATPase inhibitor EHNA (erythro-9-(2-hydroxy-3-nonyl)adenine; 10 mM), which immediately stopped the beating motion of sperm flagella^6^. After EHNA treatment, the inhibited sperm were vitrified on EM grids. To facilitate the cryoET imaging which is limited by sample thickness, lamellae of ∼300 nm-thickness were generated by cryogenic focused ion beam-scanning electron microscopy (cryo FIB-SEM). Tilt series were recorded using a 300 kV cryo transmission electron microscope (cryoTEM) and a dose-symmetric scheme with a tilting increment of 4°. The 4° tilt increment, instead of the commonly used 1-3°^7-9^, was used to improve the signal-to-noise ratio of each tilt image while keeping the same angular range and total dose. Three-dimensional classification and refinement of subtomograms corresponding to the 96 nm-repeating units were performed as reported previously (Extended Data Fig. 1)^9^. We then performed focused refinement on the 48 nm-repeating structures of doublets focusing on the microtubules, aiming to reach the highest possible resolution (see the workflow shown in Extended Data Fig. 1). The newly developed RELION4 was used to refine the 3D reconstructions and achieved 7.7 Å overall resolution (FSC = 0.143) (Fig. 1a and Extended Data Fig. 2a, b, see Methods). The improvement of resolutions compared to RELION3 comes from more accurate CTF estimation and alignment of tilt series since both parameters of each tilt image were iteratively refined relative to the 3D reconstructions (Extended Data Fig. 2a)^10^. Densities of microtubule inner proteins (MIPs) from mouse axonemes repeating every 16 nm along the axoneme axis were also observed despite of the overall periodicity of 48 nm (Fig. 1c-e). In addition, we reprocessed a previous human sperm dataset and achieved 10.3 Å for the 48 nm-repeating structures of the doublets (FSC = 0.143) (Fig. 1b and Extended Data Fig. 3a, b)^9^. Individual α-helices for the tubulins and MIPs are clearly resolved in both maps. These maps were then compared to the published cryoEM map of doublets from bovine trachea obtained by single-particle approach^11^, which were low-pass filtered to comparable resolutions of 7.5 Å and 10 Å, respectively (Extended Data Fig. 2c and 3c). Similar levels of detail in secondary and tertiary structures were resolved, validating the resolution estimates of our 3D reconstructions. To our knowledge, these resolutions are currently the highest achieved by cryoET for any *in situ* axonemal structures (12 Å maps were reported previously for equivalent structures from *Tetrahymena* cilia^8,12^).

**Figure 1.**
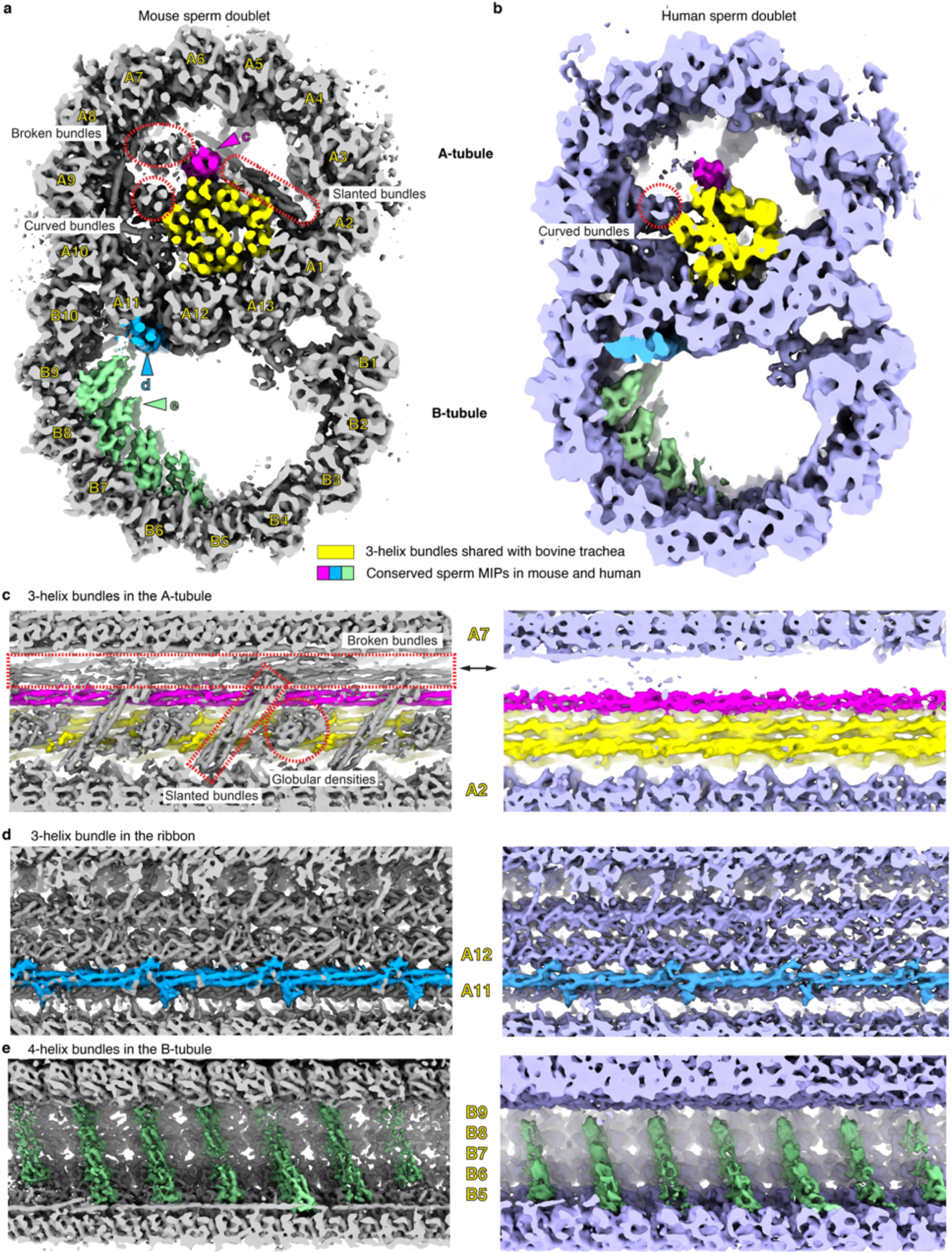
The 3D reconstructions of mouse and human sperm doublets revealed novel MIPs. **a**, Transverse cross-section views of the doublets of mouse sperm. Conserved sperm MIP densities are highlighted (pink, cyan and green) and the corresponding viewing angles of **c**-**e** are indicated (colored arrow heads). The 3-helix densities in A-tubule shared with Bovine trachea doublets (EMD-24664, REF) are colored (yellow). Divergent sperm densities are also indicated (red dashed ovals). While individual protofilaments of the doublets are labeled as A1-13 and B1-10. **b**, Transverse cross-section views of the doublets of human sperm. **c-e**. Zoom-in views of the conserved sperm MIP densities along the longitudinal axis. In **c**, mouse sperm-specific densities are indicated and labeled (see more in Extended Data Fig. 2 and 3). In **e**, although the striations are 8 nm apart, but the overall periodicity is 48 nm.

Our 3D reconstructions of mouse and human sperm doublets reveal densities similar to the map of bovine trachea doublets^11^, as well as sperm-specific densities (Fig. 1a). However, we also observed sperm-specific densities that were mostly tube-like, corresponding to bundles of α-helices. Inside the A-tubule of mouse sperm axonemes, twelve helical bundles form a filamentous core parallel to the longitudinal axis, whereas only eight helical bundles, identified as Tektin 1-4, are present in bovine trachea cilia (Fig. 1a and Extended Data Fig. 2d)^11^. Among the four mouse sperm-specific helical bundles, only one is a continuous 3-helix bundle that runs along the entire length of the doublets (Fig. 1a, c). This continuous bundle is also found in human sperm doublets (magenta densities in Fig. 1a-c). The other three bundles, the two broken bundles and the curved bundles, all have breaks within the 48-nm periodic structure and appear different in mouse and human sperm (Extended Data Fig. 2d and 3d-g). In particular, the two broken straight 3-helix bundles have very low occupancy in the human sperm doublet (Fig. 1d and Extended Data Fig. 3d, e), while one of the curved helical bundles is connected to the doublets lumen in human but not mouse sperm (Extended Data Fig. 2d and 3f, g). In the mouse sperm doublet, we also observed unique “oblique” helical densities oriented ∼45° relative to the filament axis and a globular domain next to it, both repeating every 16 nm (Fig. 1c). These comparisons suggest there are sperm-specific MIPs and also diversifications among mammalian sperm in the A-tubule of doublets.

Outside the A-tubule, we also found novel densities that are conserved in both mouse and human sperm doublets. A continuous three-helix bundle with multiple protrusions is situated at the external interface between A11 and A12 protofilaments (Fig. 1a-b, d). Inside the B-tubule, there are groups of four-helix bundles lining the inner surface of tubulins from the B4-B9 protofilaments. These four-helix bundles are stacked next to one along the helical pitch of the microtubule, consistent with the previously reported density striations at lower resolutions (Fig. 1e)^9^. Together, these data reveal that the mammalian sperm doublets have the most extensive MIP network of any microtubule structure observed so far. While the B-tubule appears similar, mouse sperm have more MIPs than human sperm in the A-tubule.

### *De novo* protein identification using AlphaFold2

The clearly resolved secondary and tertiary structures allowed us to interpret maps and to build pseudo-atomic models. First, we were able to identify densities corresponding to the 29 MIPs observed in bovine trachea cilia (Extended Data Fig. 4)^11^, suggesting their orthologs are likely present in sperm axonemes. We then sought to identify proteins contributing to the conserved sperm densities in mouse and human doublets (highlighted in Fig. 1c-e). Since most of these features repeat every 16 nm along the axoneme axis, we performed focused refinement of the 16-nm repeating structures in A- and B-tubule of mouse doublets separately and the larger number of subtomograms further improved the resolutions of averages to 6.0 Å and 6.7 Å, respectively (Extended Data Fig. 5). The unaccounted target densities from our maps were computationally isolated and unbiasedly matched to the predicted tertiary structures from the AlphaFold2 mouse proteome library (21,615 proteins) using the COLORES program from the SITUS package (Fig. 2a)^13,14^. The search time scales with the size of PDB and the volumes, and the total time for matching the mouse proteome with a volume of 60×60×60 pixels is 105 single-core hours. Dividing the 21,615 proteins for parallel runs allowed the search to be done more efficiently. The best poses for 21,615 mouse proteins were scored and ranked by the cross-correlation scores calculated by COLORES (Extended Data Fig. 6a)^14^. For the continuous 3-helix densities in the A-tubule, the best hit was Tektin 5, a Tektin found only in the mammalian testis and sperm in previous proteomic studies (Extended Data Fig. 6b-c, see methods for details)^15-17^. Although Tektin 5 has no reported structure, AlphaFold2 predicted that it adopts single-helix, 3-helix and 2-helix segments from the N- to C-termini (Fig. 2b)^13^, a tertiary structure that is almost identical to the ones reported for Tektin 1-4^11^. The ColabFold, an AlphaFold2-based Google notebook, were then used to model how two copies of Tektin 5 molecules interact to form an oligomer ^13,18^. The resulting complexes suggest the single-helix N-terminal region of one Tektin 5 could interact with the 2-helix C-termini of the other molecule (Fig. 2c), indicating its potential to self-polymerize and form a quasi-continuous 3-helix bundle. Indeed, multiple Tektin 5 could be fit into the continuous 3-helix densities with 16-nm periodicity, with little adjustment of the orientations of individual α-helices of the original AlphaFold2 model (Fig. 2d). Upon manual inspection of the hit list, Tektin 1-4 were also among the top 10 hits; this finding corroborated the robustness of our search method in finding proteins with matching tertiary structures. We assign these densities to be Tektin 5 as it is uniquely present in mammalian sperm^16,17^ and such densities are absent in bovine trachea cilia that only contain Tektin ^1-4,11^.

**Figure 2.**
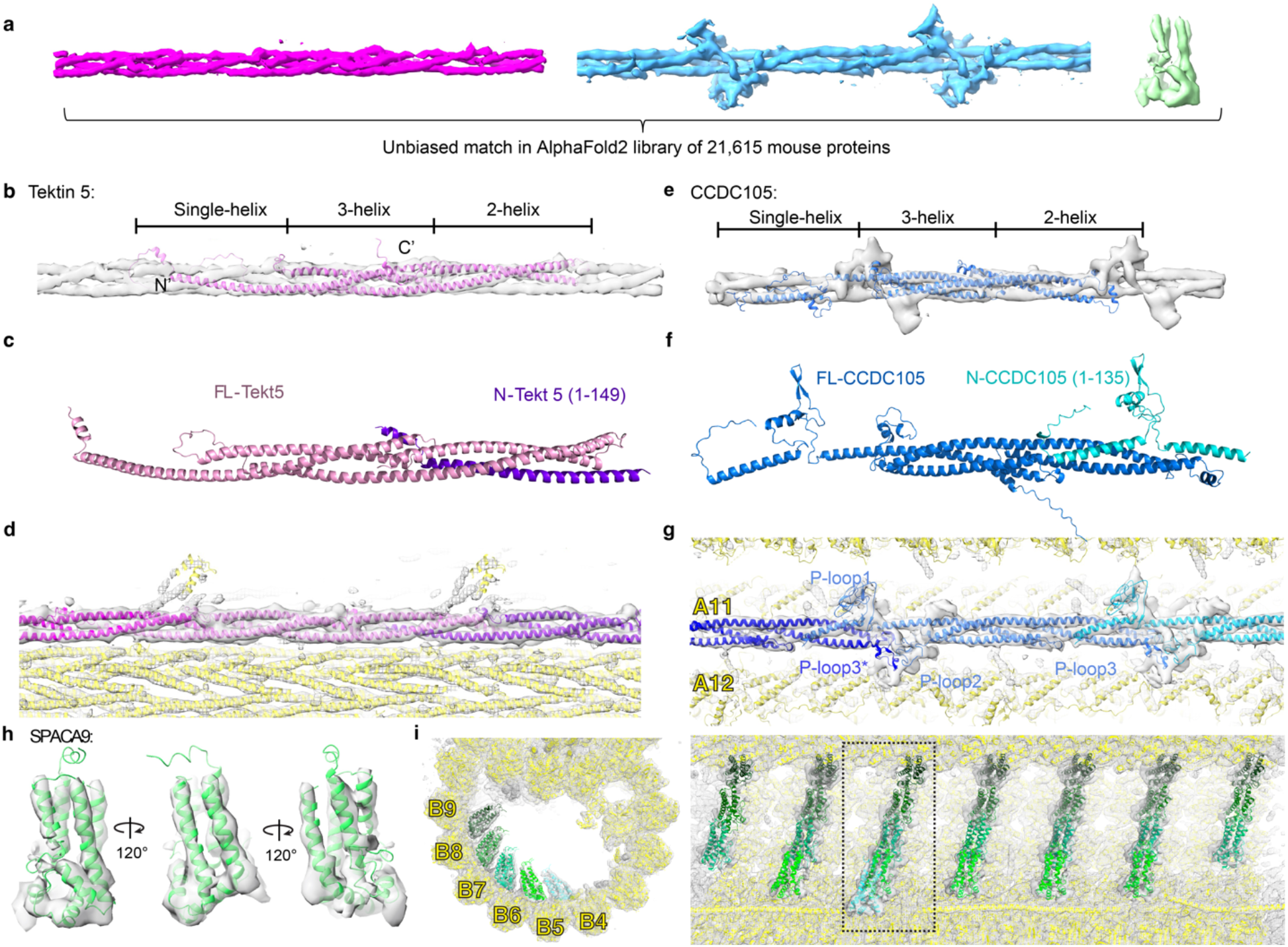
Unbiased matching of target densities from *in situ* cryoET reconstructions with structure models of 21,615 mouse proteins predicted using AlphaFold2. **a**, Unbiased search for molecules that fit into the conserved densities in mouse and human sperm. **b**, The predicted structure of Tektin 5 based on AlphaFold2. **c**. Modeling of complex formation for a full-length Tektin 5 and a truncated one. **d**, Fitting and modeling of Tektin 5 into the 3-helix bundle densities in the A-tubule. The nearby densities are accounted for by other proteins (yellow ribbon). **e**, Unbiased search in the AlphaFold2 library identified CCDC105 as the candidate for the continuous 3-helix density at the Ribbon. **f**. Two examples of modeling of complex formation for a full-length CCDC105 and a truncated one. **g**, Fitting and modeling of CCDC105 into the 3-helix bundle density at the ribbon. The nearby densities are accounted for by other proteins (yellow ribbon). **h**, The AlphaFold2 model for SPACA9 was fitted into the density directly and viewed from different angles. **i**, Two orthogonal views of the striations of SPAC9 in the B-tubule. Different SPACA9 molecules are colored with different shades of green. The left panel showed a particular striation indicated in the right panel.

For the continuous 3-helix densities with protrusions at the ribbon, CCDC105 (coiled-coil domain containing protein 105) was identified in the top 20 hits from the unbiased search in AlphaFold2 mouse proteome library, along with Tektin 1-5. CCDC105 was also found in mammalian sperm and the testis but not other tissues^15-17^. CCDC105 adopts a similar overall tertiary structure like Tektin 1-5 based on AlphaFold2 prediction (Fig. 2e)^13^, suggesting it is an uncharacterized Tektin homolog. However, there are three proline-rich loops in CCDC105 that are uniquely conserved across CCDC105 orthologs (Fig. 2e and Extended Data Fig. 6d,e) and they likely form structured loops like the ones observed in other axonemal complexes^19,20^. We also modeled how two copies of CCDC105 would interact using AlphaFold2/ColabFold^21^. The predicted interface involves coiled-coil interactions between the single-helix segment of one CCDC105 and the 2-helix segment of the other (Fig. 2f). Furthermore, CCDC105 fit well into the continuous 3-helix densities at the ribbon, with their characteristic proline-rich loops matching the protrusions in our density maps (Fig. 2g). Notably, we could not swap the fitting of Tektin 5 and CCDC105 into these two 3-helix bundles after extensive trials, mostly due to the different orientations and lengths of α-helices (Extended Data Fig. 6f).

We also extracted the 4-helix bundles at the B-tubule striations and again performed an unbiased search using the AlphaFold2 library. SPACA9 (Sperm Acrosome-Associated Protein 9) was found to be the best hit (Fig. 2h,i). SPACA9 was previously found in various ciliated organs in human (testis, fallopian tubes and lung)^15^ and its tertiary fold is so unique that no other homologous protein was found in the top-200 ranked structures from the unbiased search. Interestingly, no match was found when the search was done against the CATH library that curates non-redundant domains of published PDBs^22^. Thus, the capability of AlphaFold2 to predict protein structure accurately, especially for the ones without published homologous structures, is critical to carrying out the unbiased proteome-wide survey.

We next examined the mouse sperm-specific densities. The globular density in the A-tubule that repeats every 16 nm was found to match the tertiary structure of DUSP proteins (Dual Specificity Phosphatase) (Fig. 3c and Extended Data Fig. 7). There are also several 3-helix bundles appear to be similar to the ones formed by Tektins 1-5, apart from the discontinuous sections and do not form continuous filaments (Fig. 3a and Extended Data Fig. 2d). We also applied the unbiased search method to the slanted and curved helical bundles (Fig. 1c and Extended Data Fig. 2e). Intriguingly, Tektin 1-5 and CCDC105 were found to be among the top 30 hits in both cases while no other PDB among the top 200 fits better, albeit only part of the structures are observed for by the densities. For the slanted helical densities, there is an additional α-helix connecting to the position where the missing single helix was expected to stem from and is folded back by ∼180° (Fig. 3b,c). Interestingly, Tektin 5, but not Tektin 1-4, has multiple conserved Gly residues among its orthologs at this turning region, so it is possible that Tektin 5 could adopt a bent-helix conformation. For the curved helical bundles, there are three 16-nm groups of densities within every 48 nm repeats (Fig. 3d and Extended Data Fig. 2d). These densities could be explained by three modified Tektin 5 molecules, in which two out of the three oligomerization interfaces near the two NME7 (a previously described MIP in bovine trachea cilia) are disrupted (compare Fig. 3d with Fig. 2c,d). The first and second Tektin 5s are missing the single-helix segment beyond the conserved Gly137 (mouse), while the second and third Tektin 5s have curved 2-helix segments. These modifications of the AlphaFold2-predicted Tektin 5 are necessary to avoid spatial clash with the two NME7 (Fig. 3b and Extended Data Fig. 4b). As these structures were not observed in bovine trachea cilia that only contains Tektin 1-4^11^, we hypothesize that Tektin 5 has evolved to adopt multiple conformations and positions within sperm axonemes.

**Figure 3.**
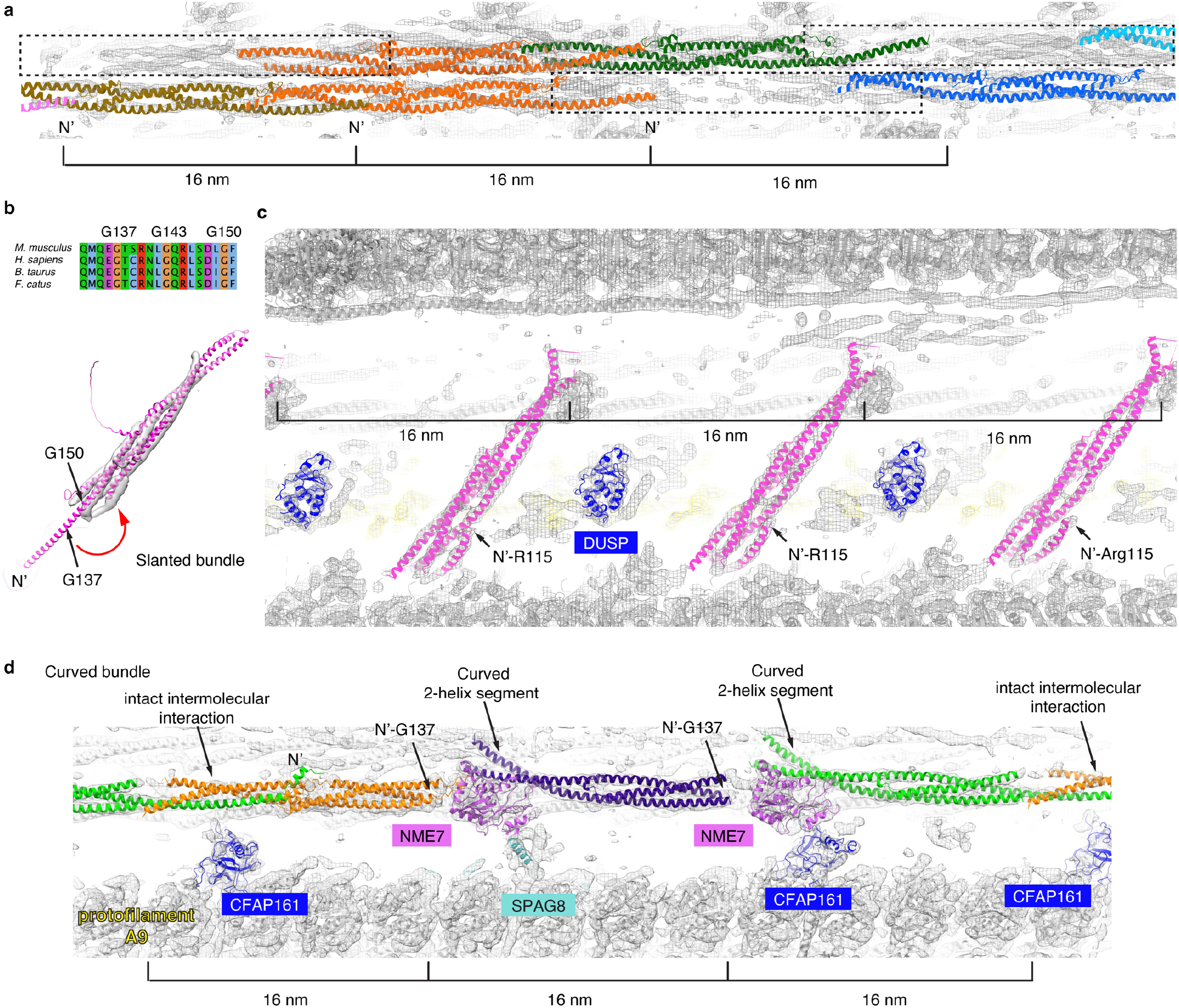
Protein candidates for mouse sperm-specific densities and conformational plasticity of Tektin 5. **a**, The two broken 3-helix bundles could be explained by two complete and a third partial copies of Tektin 5 per 48-nm repeat, instead of three in the continuous 3-helix bundle. **b**, The AlphaFold2 model of mouse Tektin 5 was fit into the slanted helical densities in the reconstruction of 16 nm-repeating doublet structure. Sequence alignment of Tektin 5 from mouse, human, bovine and cat is shown from Q133-F151 (the numbers are based on *Mus musculus* Tektin 5). Conserved G137, G143 and G150 are near the turning point of the bent α-helix. **c**, The fitting of Tektin 5 and DUSP3 protein into the 16-nm repeating features, see the same view of the map in Fig. 1c. **d**, Three modified Tektin 5 were fitted into the densities of curved bundles in the mouse sperm doublet (Fig. 1a). The presence of two NME7 correlate with disrupted the inter-molecular interfaces of Tektin 5 shown in Fig. 2c.

### Orthogonal proteomic analyses

To further validate our *de novo* protein assignments based upon AlphaFold2 modeling and docking to EM maps from cellular cryoET, we used mass spectrometry-based proteomics. Mouse sperm were isolated and extracted using salt buffers with increasing denaturing capabilities (E1: 0.1% Triton, E2: NaCl, E3: KSCN, E4: Urea, and E5: 10% SDS) (Extended Data Fig. 8a). α-tubulins could be detected in KSCN and Urea extractions but not in others (Extended Data Fig. 8b), suggesting the microtubule doublets are disassembled and the MIP candidates are likely present in these two extractions. After analyzing the E1-E5 fractions by MS, proteins with significant changes in abundance between fractions were clustered into six distinct groups based on correlation of intensity profile (Extended Data Fig. 8c,d and Supplementary Table S1,2)^23,24^. Gene ontology (GO) analyses suggest that cluster 4 is enriched for proteins involved in cilium, cilium assembly, cytoskeleton and the axoneme, and shows increased intensities in fraction E3, while cluster 6 is enriched for cilia assembly and shows increased intensities in fraction E4 (Extended Data Fig. 8e and Supplementary Table S3)^25^. The overall protein abundance is consistent with the idea that E3 and E4 buffers extracted axonemal proteins. Indeed, 28 of the 29 previously identified MIPs in bovine trachea doublets were reproducibly identified in all three biological replicates (Supplementary Table S4). The almost complete list of MIPs highlighted the coverage of our biochemical and MS analyses. Importantly, SPACA9 was reproducibly identified in fraction E3, while Tektin 1-5 and CCDC105 were reproducibly identified in fractions E3 and E4, with high protein intensity and a range of 3 to 34 unique peptide identifications per replicate (Extended Data Fig. 8f, Supplementary Table S4). Only one DUSP protein, DUSP3, was identified in the in two of three replicates in fraction E4 fraction (Extended Data Fig. 8f, Supplementary Table S4). Moreover, AlphaFold2 models for the other candidates from fractions E1-4 were inspected but no additional candidates could explain the characteristic helical densities described above.

## Discussion

Our *in situ* cryoET studies have allowed high-resolution imaging of the structures and interactions of macromolecular complexes in mouse and human sperm. The well-resolved secondary and tertiary structures combined with an unbiased search using the library of AlphaFold2-predicted models of the mouse proteome allowed us to identify Tektin 5, SPACA9 and CCDC105 in sperm doublets based on well-matched and unique fittings; Genetic knock-out experiments followed by *in situ* cryoET could provide further verification of these assignments.

Our reconstructions revealed an extensive interaction network inside the sperm doublets formed by the novel proteins. In the sperm doublets, SPACA9 and CCDC105 directly crosslink the tubulin dimers longitudinally and laterally at unoccupied lumen sites of bovine trachea doublet microtubules but do not affect the positions of “common MIPs” (those shared by mammalian tracheal cilia and sperm flagella) (Fig. 2 g,I and Extended Data Fig. 4)^11^. Tektin 5 adopts various conformations around the filamentous core formed by Tektin 1-4. The conformational plasticity observed for Tektin 5 is partially shaped by the limited space, as shown by the bent single helix that would otherwise clash with the microtubule wall (Fig. 3b,c), as well as the missing single helix and curved 2-helix segment near NME7 (Fig. 3d). These observations are consistent with a stepwise assembling model where MIPs shared by mammalian trachea cilia and sperm flagella assemble first and SPACA9, CCDC105 and non-polymerizing Tektin 5 are recruited later.

In sperm A-tubules, the 3-helix bundles of Tektin 1-5 are arranged with different polarities (Fig. 4a) and orientations (Fig. 4b). We hypothesize that this arrangement may allow the outer doublet to withstand additional mechanical stress from different directions as the 3D bending pattern of mammalian sperm would stress the nine doublets from different angles (Fig. 4c)^26^. The intramolecular and intermolecular coiled-coil interaction interfaces are parallel to the microtubule axis so that bending would not expand the gap of these interfaces (Fig. 4d). Instead, the bending force would lead to bent helical bundles and distort the ideal bond angles/lengths. The transition of releasing such molecular strains could provide a restoring force that allows a curved filament to return to a straight conformation. In contrast, the interface between tubulin dimers along the protofilaments is in a plane perpendicular to the filament axis; bending would open the interface and lower the affinity or the potential restoring force.

**Figure 4.**
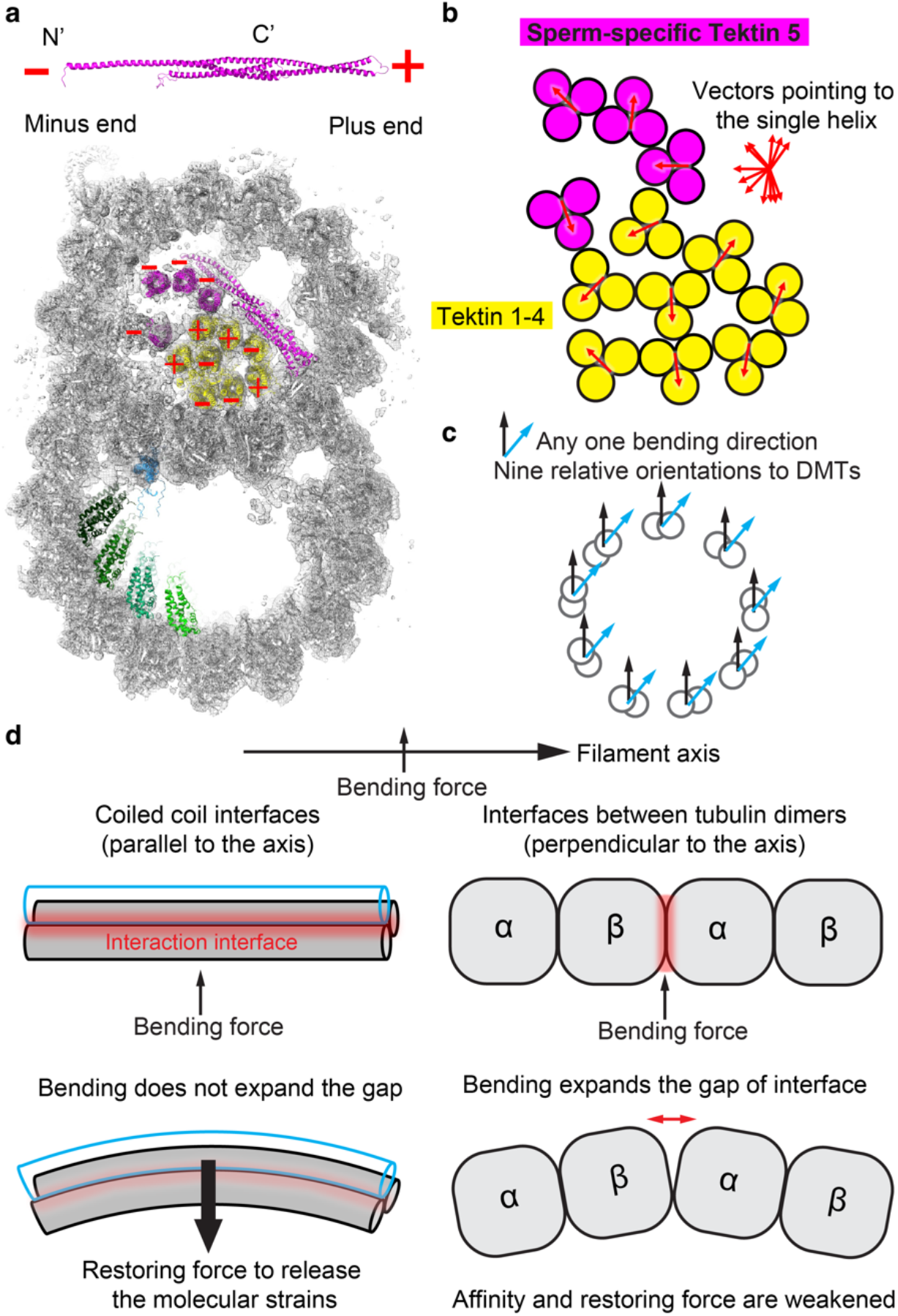
Sperm doublets are composed of microtubules and intermediate filaments. **a**, The plus and minus ends of Tektin 5 were named based on the N- and C-termini of the protein. The cross-section view of the doublets showing the polarities of 3-helix bundles pointing towards the readers. **b**, The orientations for each 3-helix bundle were represented by a vector starting from the middle point of the 2-helix segment and pointing to the single-helix segment of the other molecule. **c**, In the context of the (9+2) axoneme, bending would lead to stress of the nine doublets from nine directions. Two different directions of bending are shown (black and blue arrows). **d**, A model of how 3-helix bundles could improve the mechanical stability and elasticity of the microtubules. The bending curvature and gap are exaggerated for illustration purposes.

Here, we describe a workflow of combining high-resolution *in situ* cryoET and AlphaFold2 modeling that defines a visual proteomics approach of precisely placing proteins in their native cellular environment without labeling, cellular disruption and purification. This workflow has allowed us to uncover the cellular locations and interaction networks of MIPs, which suggests hypothesis of how they contribute to the mechanics of flagellar bending. This visual proteomic workflow could potentially be applied to other cell biology problems, such as membrane remodeling by viruses and identification of their *in situ* interactors.

---

**Extended Data Fig. 1.**
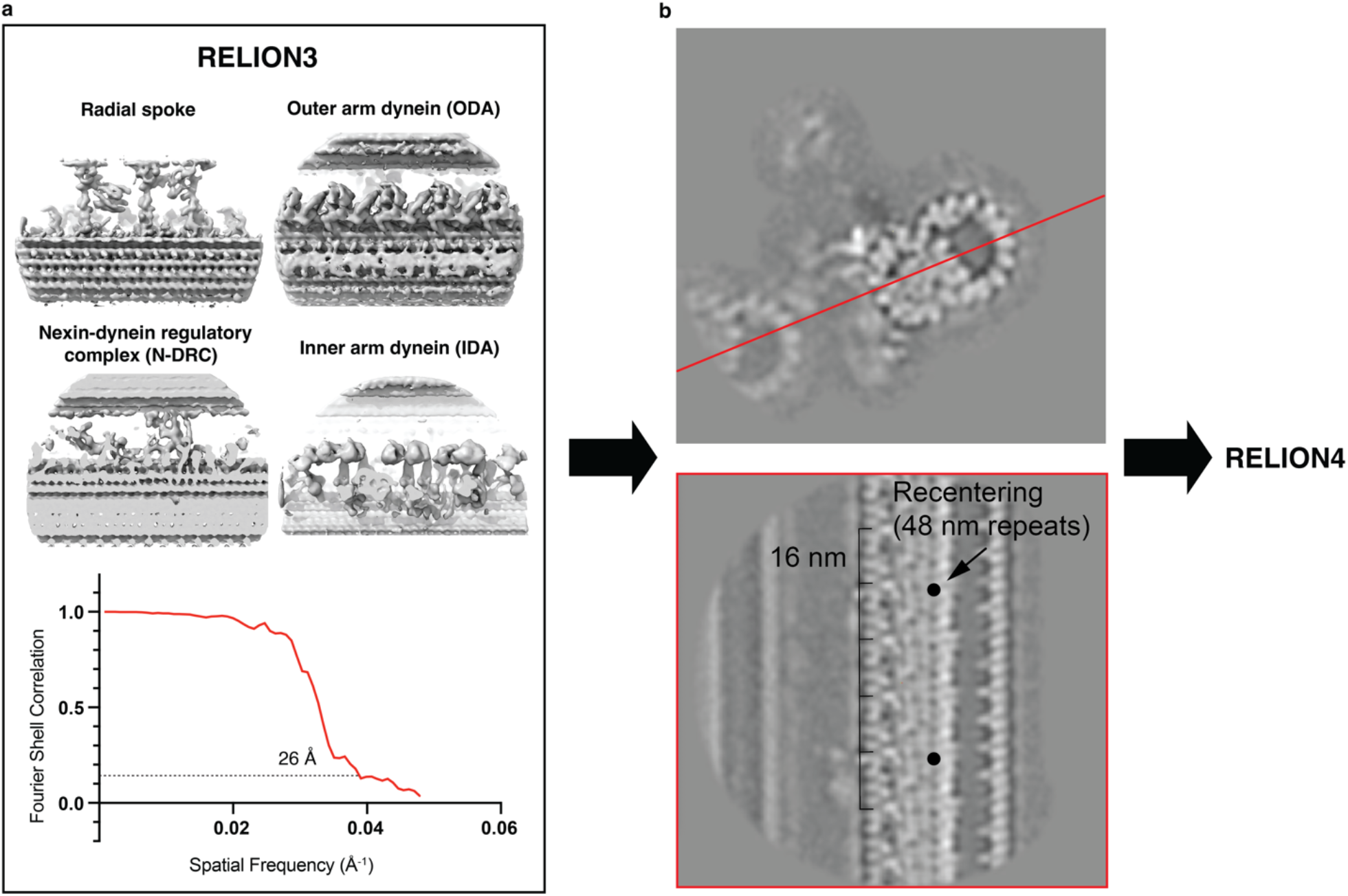
Workflow of data processing. **a**, Four views of the 96 nm-repeating structure of doublets from EHNA-treated are shown for the 3D reconstruction generated using RELION3. Gold-standard Fourier Shell Correlation (FSC) curve calculated between half maps of mouse sperm doublets. The resolution was estimated as 26 Å (FSC = 0.143). **b** Two slices of the 96 nm-repeating structure of doublets looking along and perpendicular to the filament axis. Note the red line in the top panel indicates the plane of bottom slice and periodic structures are observed inside the microtubules. The coordinates were recentered on the 48-nm repeats and imported into RELION4. In the top panel, note the features further away from the microtubules are more blurry, suggesting that there are conformational heterogeneities and they are resolved at lower resolutions.

**Extended Data Fig. 2.**
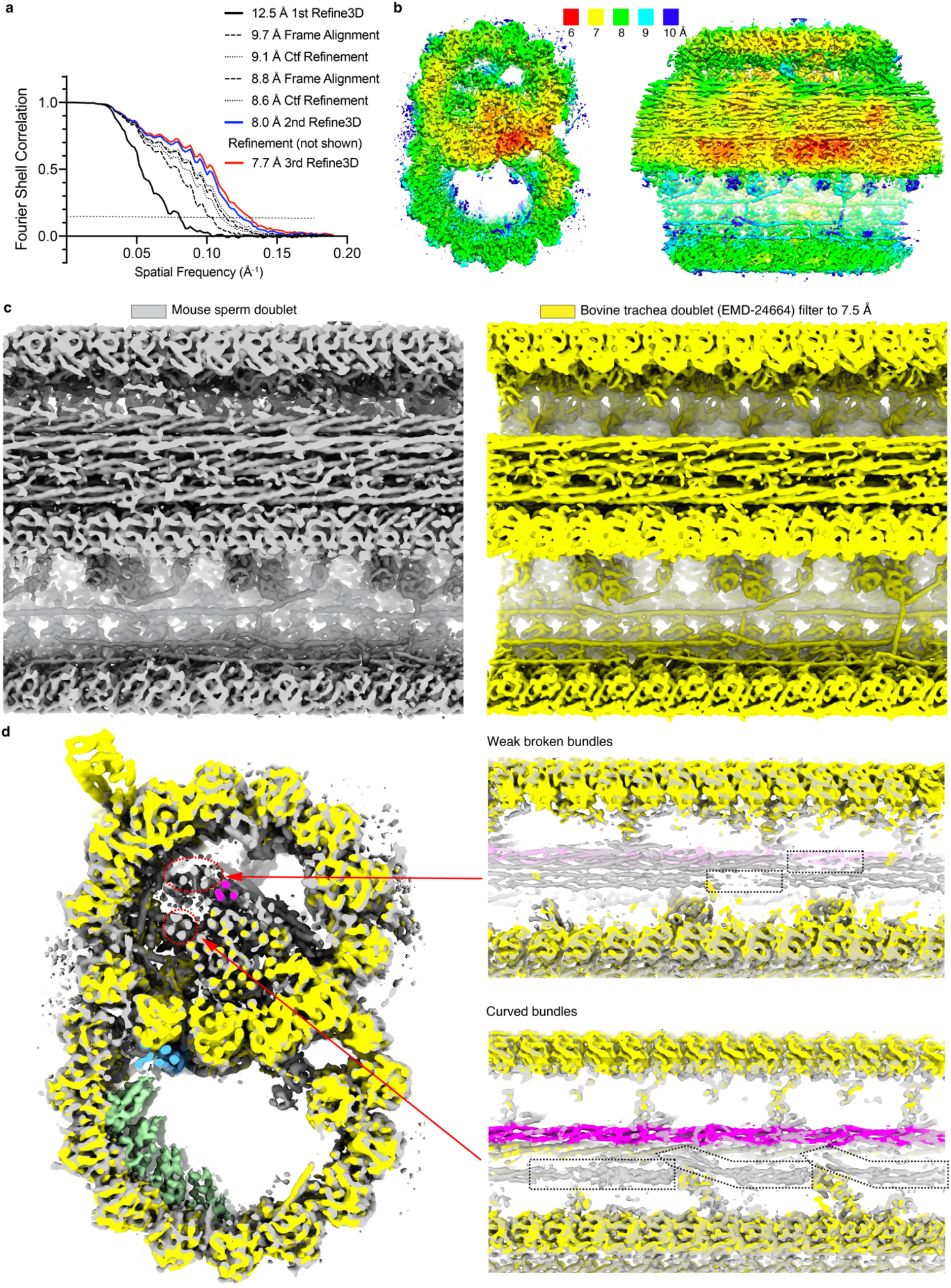
Characterization of the 48 nm-repeating structure of doublets from mouse sperm. **a**, Gold-standard Fourier Shell Correlation (FSC) curves calculated between half maps of mouse sperm doublets. The resolutions were reported as FSC = 0.143. Note the FSC curves resulted from the iterative frame alignment and ctf refinement between the second and third Refine3D jobs were not shown for the clarity of the figure. Further refinement did not improve the resolution or quality of the map. **b**, Local-resolution maps of mouse sperm doublets calculated by RELION4. The ribbon region has the highest resolutions. Densities in the A-tubule have higher resolutions than the ones from the B-tubule. **c**, Equivalent longitudinal cross-section views of doublets from mouse sperm and bovine trachea cilia (EMD-24664) are shown. The latter was low-pass filtered to 7.5 Å and comparable details of the secondary and tertiary structures of the MIPs are observed. **d**, Zoom-in views of the two broken helical bundles and the curved helical bundles inside the A-tubule of mouse sperm doublets. The discontinuous parts of the broken helical bundles are indicated (dashed shapes). Note the curved bundles have a straight and two curved groups of densities (outlined using dashed shapes).

**Extended Data Fig. 3.**
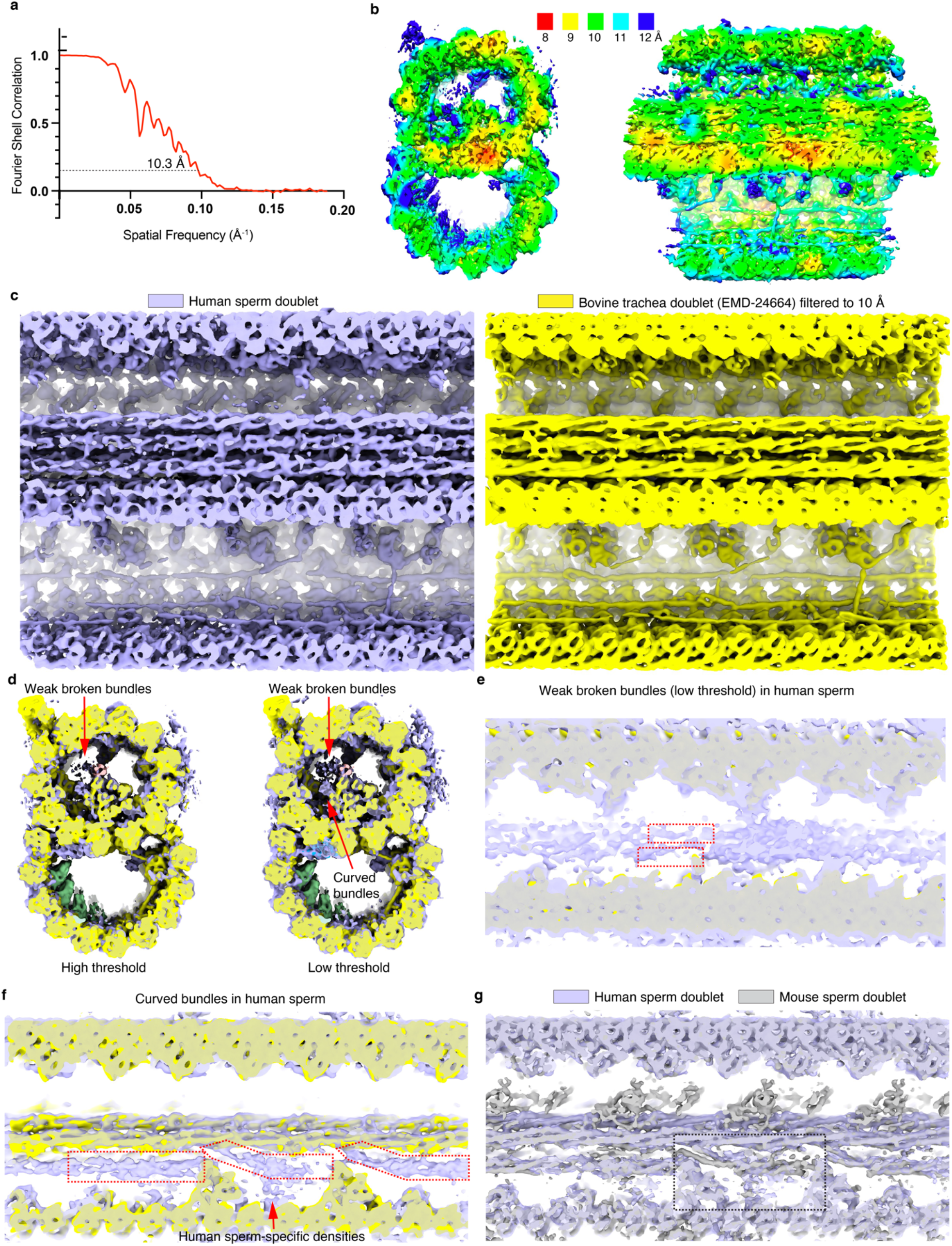
Characterization of the 48 nm-repeating structure of doublets from human sperm. **a**, Gold-standard Fourier Shell Correlation (FSC) curve calculated between half maps of mouse sperm doublets. The resolution was estimated as 10.3 Å (FSC = 0.143). **b**, Local-resolution maps of human sperm doublets calculated by RELION4. The ribbon region has the highest resolutions. Densities in the A-tubule have higher resolutions than the ones from the B-tubule. **c**, Equivalent views of doublets from human sperm and bovine trachea cilia (EMD-24664, REF) are shown. The latter was low-pass filtered to 10 Å and comparable details of the secondary and tertiary structures of the MIPs are observed. **d**, Two cross-section views of the human sperm doublets overlaid with bovine trachea doublets (EMD-24664) at low and high thresholds. **e**, The two broken bundles inside the A-tubule in human sperm are shown at low threshold (see the corresponding mouse densities in Extended Data Fig. 2d). **f**, The curved helical bundles contain a straight and two curved groups of densities inside the A-tubule of human sperm are outlined. Human sperm-specific densities were observed to connect one curved bundles to the lumen of A-tubule. **g**, The human sperm doublets overlaid with mouse sperm doublets are shown. The inconsistent densities are outlined (dashed line) (also see Extended Data Fig. 2d and 3f).

**Extended Data Fig. 4.**
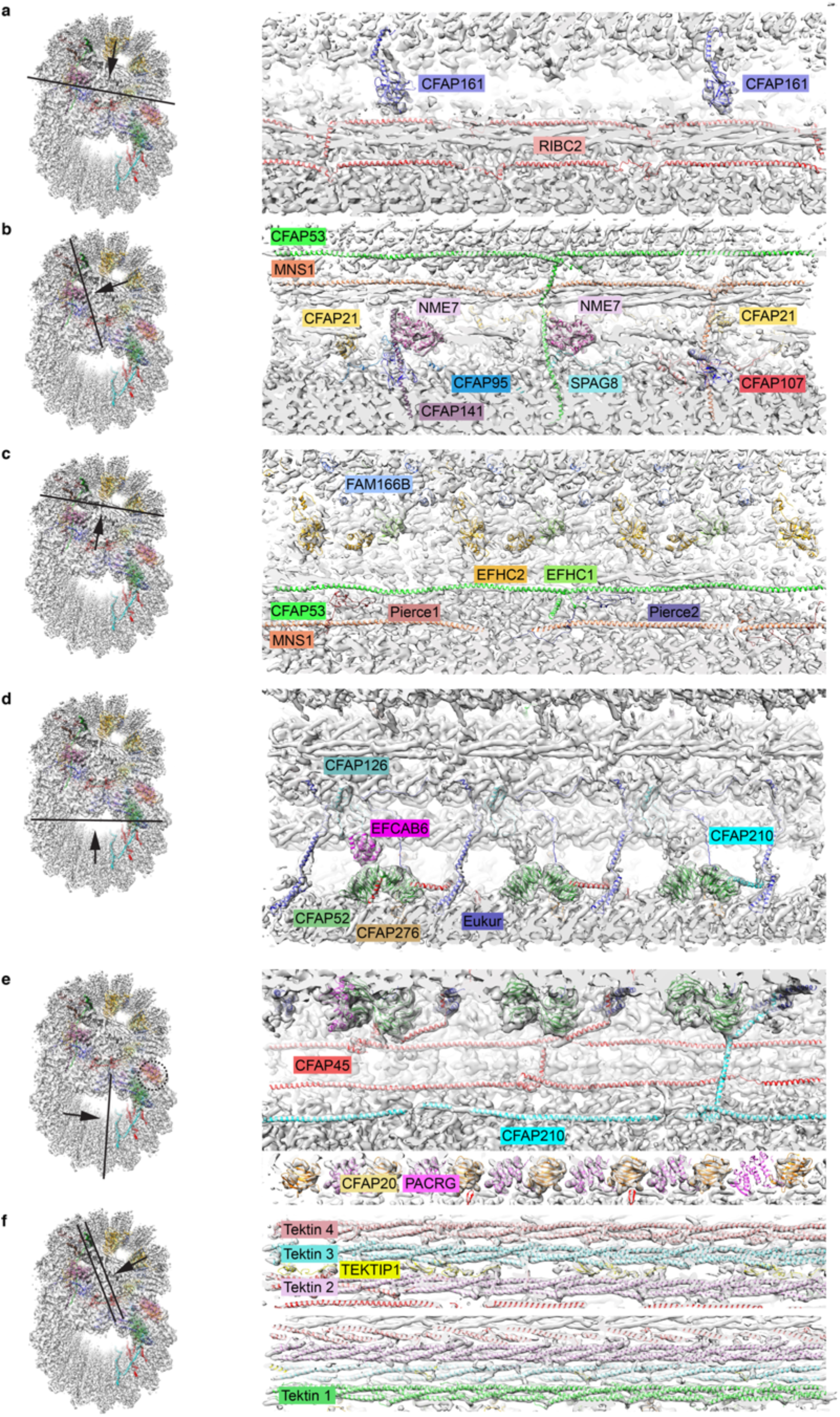
Rigid-body fitting of 29 known MIPs from bovine trachea cilia into the density map of mouse sperm doublet. **-f**, Models of 29 known MIPs from bovine trachea cilia (PDB 7rro, REF) are fitted into the density map of mouse sperm doublet. The viewing angles for all panels are shown. For proteins that have multiple α-helices (CFAP161, RIBC2, CFAP53, MNS1, CFAP21, NME7, CFAP141, EFHC1, EFHC2, ENKUR, CFAP210, EFCAB6, CFAP45, PACRG and TEKTIN 1-4), the arrangement of secondary structures matches densities in sperm doublets. The overall shapes of Beta-sheet-rich proteins (CFAP52 and CFAP20) match the densities and these proteins are highly conserved in axonemes. For the proteins that contain random coils, we did observe matching features in the maps but it is generally harder to trace the main chains at the current resolution (CFAP95, SPAG8, CFAP107, FAM166B, Pierce1, Pierce2, CFAP126, CFAP276 and TEKTIP1).

**Extended Data Fig. 5.**
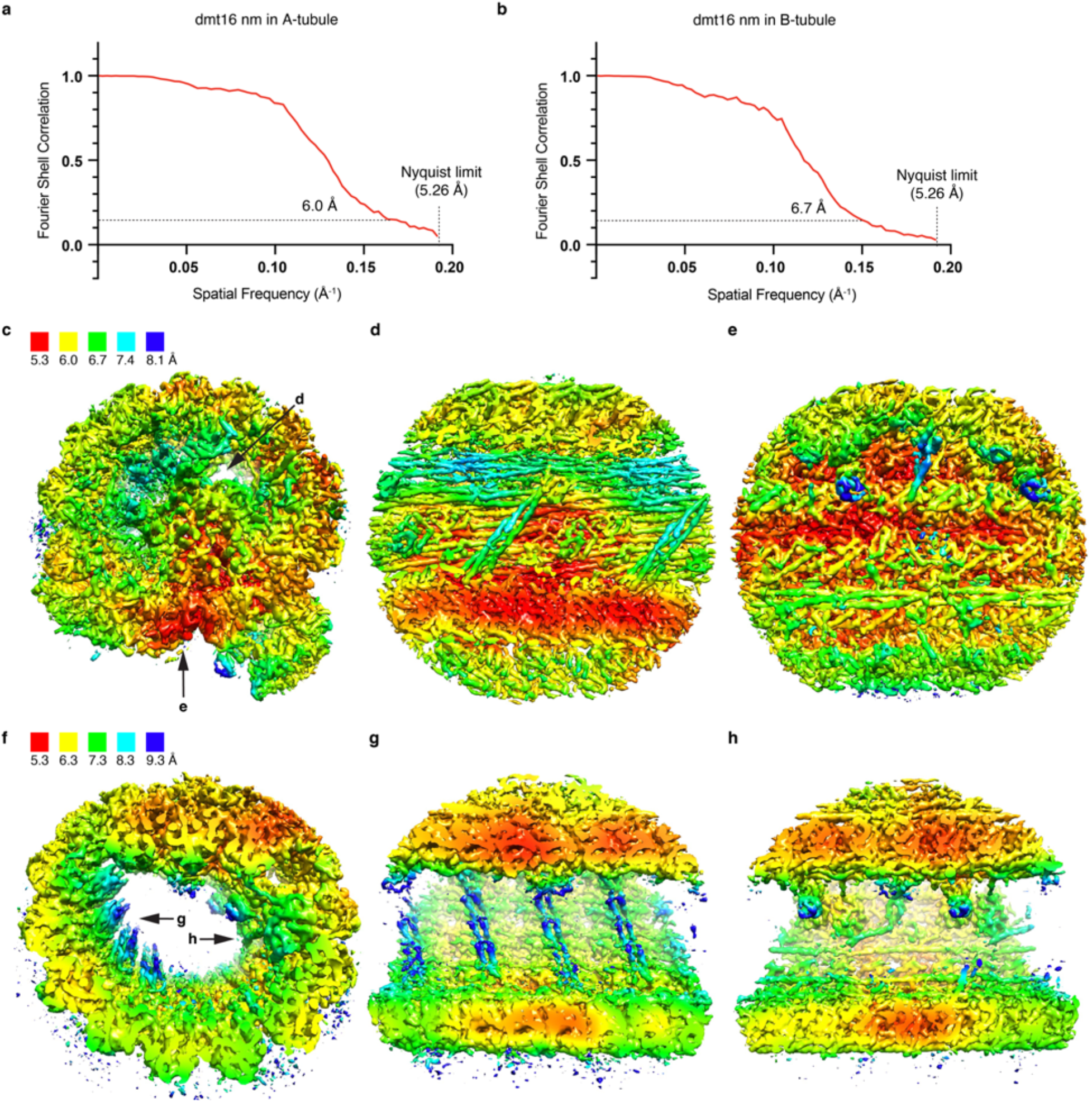
Characterization of the 16 nm-repeating structures of doublets from mouse sperm. **a**,**b**, Gold-standard Fourier Shell Correlation (FSC) curve and local resolution maps calculated using half maps of 16 nm-repeating structures of A-tubule and B-tubule. The resolution was estimated as 6.0 Å and 6.7 Å, respectively (FSC = 0.143). The Nyquist limit is 5.26 Å. **c-e**, Three views of local resolution maps calculated using half maps of 16 nm-repeating structures of A-tubule using RELION4. The viewing angles for **d** and **e** are shown in **c** (black arrow). These viewing angles are similar to Fig. 1a, d and e, respectively. Note the turns of α-helices are observed as bumps instead of a smooth cylindric density. **f-h**, Three views of local resolution maps calculated using half maps of 16 nm-repeating structures of B-tubule using RELION4. The viewing angles for **g** and **h** are shown in **f** (black arrow). The viewing angles of f and g are similar to Fig. 1a and f, respectively.

**Extended Data Fig. 6.**
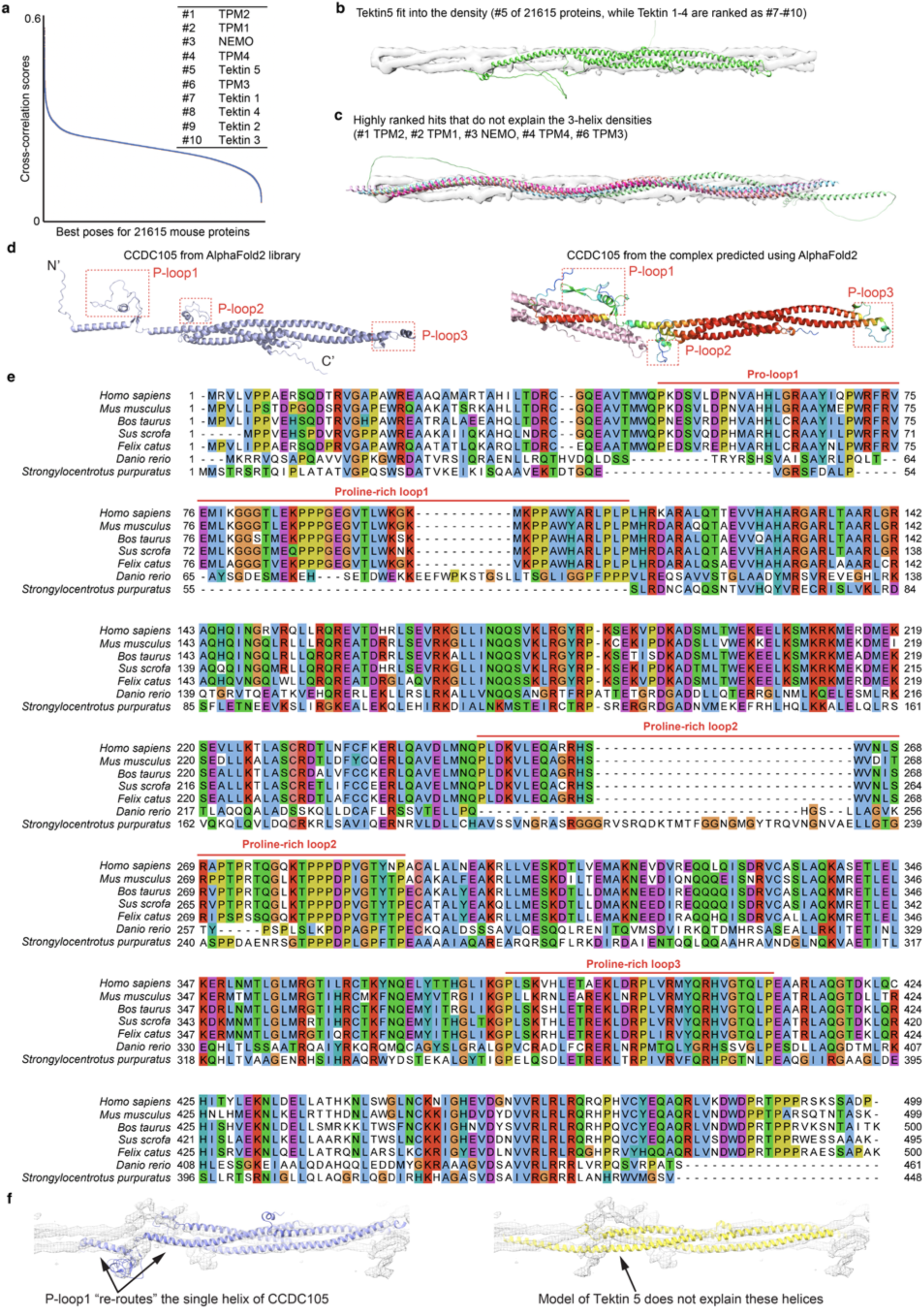
Tektin 5 and CCDC105 likely form sperm-specific 3-helix bundles associated with the A-tubule. **a**,**b**, Tektin 5 was scored as the #5 hit of the predicted structures of 21615 proteins from the mouse proteome, ranked by cross-correlation scores (Top 10 are shown). Tektin 1-4 were ranked at #7-10 due to their similar tertiary structures. **c**, Typical false positives (#1-4 and #6) from the same search. Usually these are proteins with a long single helix that matches the length of the densities but they do not explain the 3-helix densities. **d**, The structure of CCDC105 predicted by AlphaFold2 directly (left) is compared to the predicted complex formed by two CCDC105 molecules (right). The full-length CCDC105 molecule in the complex is colored based on the per-residue confidence scores (predicted local distance difference test, or pLDDT) from the AlphaFold2 prediction. The three P-loops have medium confidence scores (green), suggesting the exact conformations of these loops may not be accurately predicted. However, the presence of these structured loops is conceivably confident based on the conserved proline residues (see the sequence alignment in **f**) and matched the protrusion densities observed in our maps (Fig. 2g). Note the conformations of the three proline-rich loops differ in these two predictions. These differences could be caused by the presence of neighboring molecules (P-loop1 and P-loop2) or the different AlphaFold2 workflows (REFs). **e**, The sequence alignment of CCDC105 from five mammals (*Homo sapiens, Mus musculus, Bos taurus, Sus scrofa* and *Felix catus*), zebrafish (*Danio rerio*) and sea urchins (*Strongylocentrotus purpuratus*). The proline-rich loops are marked above the sequences. **f**, The models of CCDC105 and Tektin 5 are fitted into the densities of the 3-helix bundle at the ribbon, where the former model explains the extra protrusions and orientation/lengths of helices of the densities but the latter does not.

**Extended Data Fig. 7.**
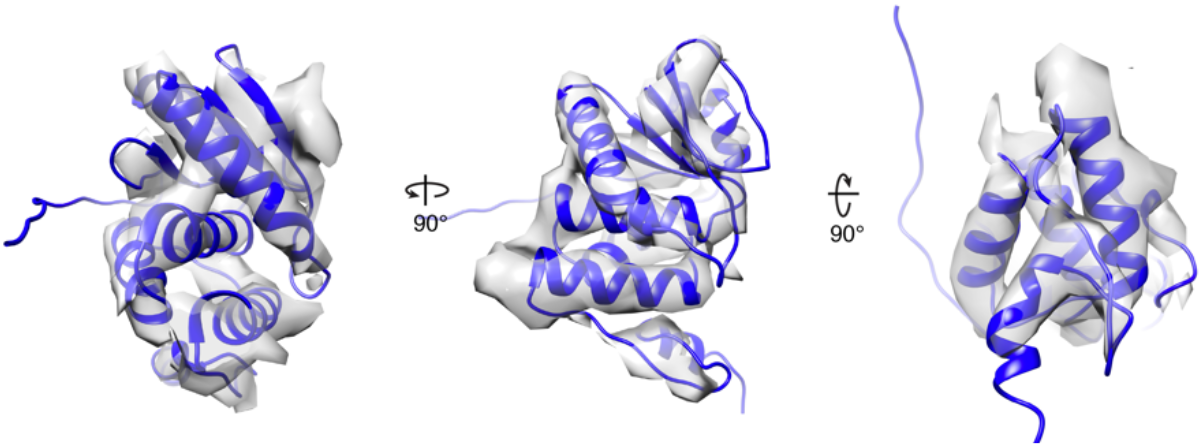
Fitting of DUSP protein. The DUSP3 is fitted into the globular domain and three orthogonal views are shown.

**Extended Data Fig. 8.**
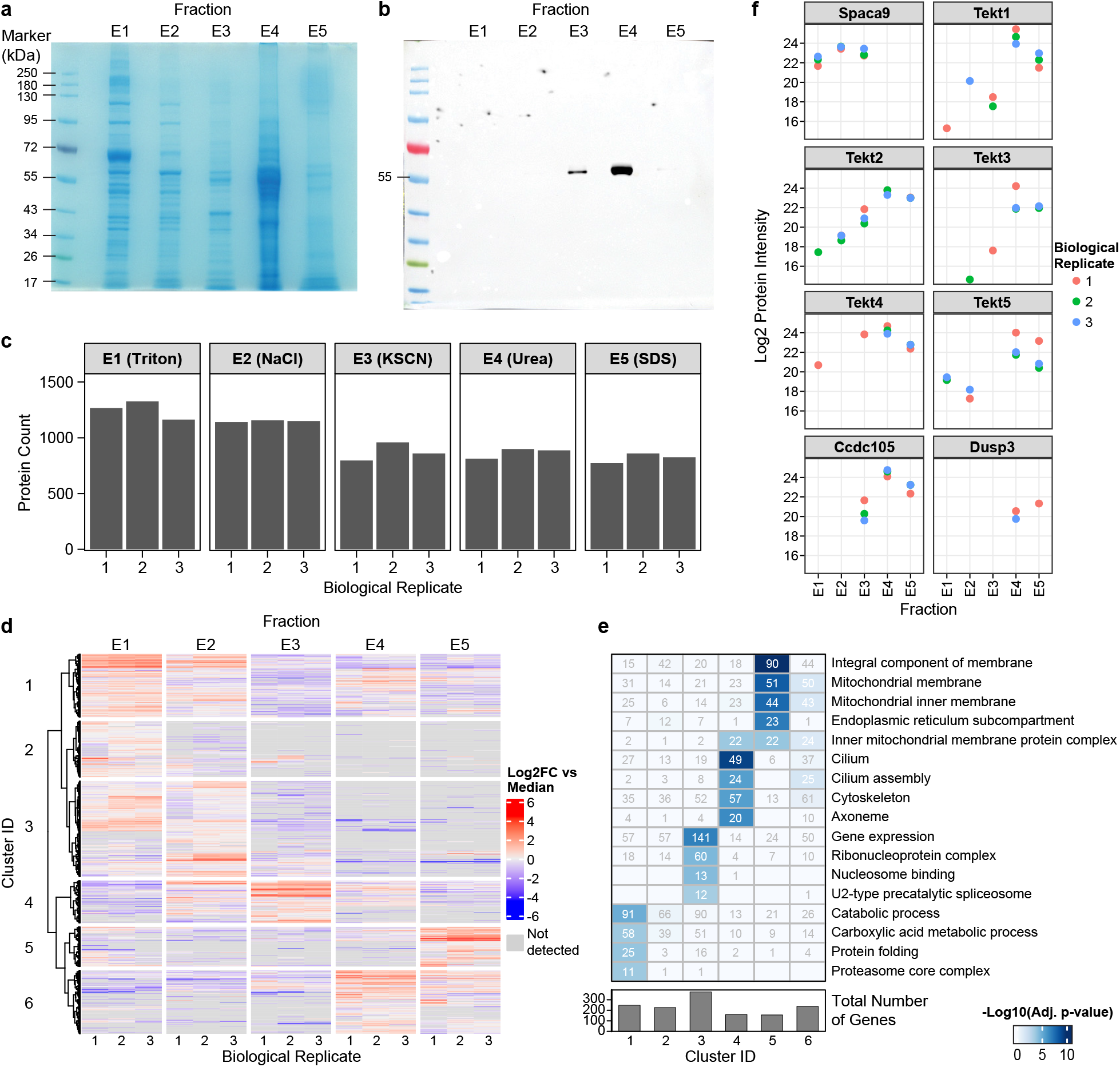
Biochemical extractions of proteins from mouse sperm. **a**, SDS-PAGE analyses of protein extractions from mouse sperm using 0.1 % Triton in PBS (E1), 0.6 M NaCl in PBS (E2), 0.6 M KCSN in PBS (E3), 8 M urea (E4) and 10% SDS (E5). **b**, Western blot analyses of protein extractions from mouse sperm using antibody against α-tubulins. Note strong bands were detected only in E3 and E4, suggesting the microtubule structures were stable in Triton and high NaCl buffer, and dissembled completely in KCSN/urea solutions.

## Data availability

MS data are shared and available through the ProteomeXchange Consortium via the PRIDE partner repository under the dataset identifier: PXD036885 (username; password: tMEZ90MC)^27^.

## Code availability

R package source materials for MSstats (version 3) are publicly available through the Krogan Lab GitHub: https://github.com/kroganlab.

## Author contributions

Z.C., R.D.V. and D.A.A. conceived the project and designed the experiments with input from other authors. S.Z. isolated the mouse sperm sample. M.S. prepared mouse sperm grids and performed the FIB milling. Z.C. collected and processed the cryoET data with suggestions from D.A.A.. Z.C. performed the biochemical extraction of mouse sperm. K.M.H. and B.J.P. acquired and analyzed the mass spectrometry data, with input and supervision from N.J.K. and R.M.K. Z.C., K.M.H., R. M.K., R.D.V. and D.A.A. wrote the manuscript draft with comments from all authors.

## Acknowledgement

We are grateful to members of the Agard and Vale laboratories for discussions and critical reading of the manuscript. We thank Xiaowei Zhao, Shixin Yang and Rui Yan from the CryoEM facility at Janelia Research Campus for their assistance with data collection. We thank Zanlin Yu for his suggestions on sample processing and model building. We thank Garrett Greenan, Shawn Zheng and Sam Li at UCSF for discussions on cryoET data processing. EM data processing utilized computing resources at both the workstations at the cryoEM facility at Janelia Research Campus and UCSF HPC Wynton cluster. We are grateful to members of the Vale laboratory and Agard laboratory for discussions and critical reading of the manuscript. Z.C. was supported by the Helen Hay Whitney Foundation Postdoctoral Fellowship. D.A.A. received funding from NIH R35GM118099. R.D.V. received funding from NIH R35GM118106 and the Howard Hughes Medical Institute. We also thank David Bulkley, Glenn Gilbert, and Matt Harrington from the UCSF cryoEM facility for the discussions on data collection and processing and NIH instrumentation grants 1S10OD026881, 1S10OD020054, and 1S10OD021741.

## Reference

1. Sironen, A., Shoemark, A., Patel, M., Loebinger, M.R. & Mitchison, H.M. Sperm defects in primary ciliary dyskinesia and related causes of male infertility. Cell Mol Life Sci 77, 2029–2048 (2020).

2. Ishikawa, T. Axoneme Structure from Motile Cilia. Cold Spring Harb Perspect Biol 9(2017).

3. Fawcett, D.W. The mammalian spermatozoon. Dev Biol 44, 394–436 (1975).

4. Lindemann, C.B. Functional significance of the outer dense fibers of mammalian sperm examined by computer simulations with the geometric clutch model. Cell Motil Cytoskeleton 34, 258–70 (1996).

5. Lindemann, C.B. & Lesich, K.A. Functional anatomy of the mammalian sperm flagellum. Cytoskeleton (Hoboken) 73, 652–669 (2016).

6. Bouchard, P., Penningroth, S.M., Cheung, A., Gagnon, C. & Bardin, C.W. erythro-9-[3-(2-Hydroxynonyl)]adenine is an inhibitor of sperm motility that blocks dynein ATPase and protein carboxylmethylase activities. Proc Natl Acad Sci U S A 78, 1033–6 (1981).

7. Nicastro, D. et al. The molecular architecture of axonemes revealed by cryoelectron tomography. Science 313, 944–8 (2006).

8. Li, S., Fernandez, J.J., Fabritius, A.S., Agard, D.A. & Winey, M. Electron cryo-tomography structure of axonemal doublet microtubule from Tetrahymena thermophila. Life Sci Alliance 5(2022).

9. Chen, Z. et al. In situ cryo-electron tomography reveals the asymmetric architecture of mammalian sperm axonemes.. Nat Struct Mol Biol, in press (2022).

10. Zivanov, J. et al. A Bayesian approach to single-particle electron cryo-tomography in RELION-4.0. Biorxiv, https://www.biorxiv.org/content/10.1101/2022.02.28.482229v1 (2022).

11. Gui, M. et al. De novo identification of mammalian ciliary motility proteins using cryo-EM. Cell 184, 5791–5806 e19 (2021).

12. Song, K. et al. In situ structure determination at nanometer resolution using TYGRESS. Nat Methods 17, 201–208 (2020).

13. Jumper, J. et al. Highly accurate protein structure prediction with AlphaFold. Nature 596, 583–589 (2021).

14. Wriggers, W., Milligan, R.A. & McCammon, J.A. Situs: A package for docking crystal structures into low-resolution maps from electron microscopy. J Struct Biol 125, 185–95 (1999).

15. Uhlen, M. et al. Proteomics. Tissue-based map of the human proteome. Science 347, 1260419 (2015).

16. Baker, M.A., Hetherington, L., Reeves, G.M. & Aitken, R.J. The mouse sperm proteome characterized via IPG strip prefractionation and LC-MS/MS identification. Proteomics 8, 1720–30 (2008).

17. Firat-Karalar, E.N., Sante, J., Elliott, S. & Stearns, T. Proteomic analysis of mammalian sperm cells identifies new components of the centrosome. J Cell Sci 127, 4128–33 (2014).

18. Madeira, F. et al. Search and sequence analysis tools services from EMBL-EBI in 2022. Nucleic Acids Res (2022).

19. Han, L. et al. Cryo-EM structure of an active central apparatus. Nat Struct Mol Biol 29, 472–482 (2022).

20. Rupp, G., O’Toole, E. & Porter, M.E. The Chlamydomonas PF6 locus encodes a large alanine/proline-rich polypeptide that is required for assembly of a central pair projection and regulates flagellar motility. Mol Biol Cell 12, 739–51 (2001).

21. Mirdita, M. et al. ColabFold: making protein folding accessible to all. Nat Methods 19, 679–682 (2022).

22. Sillitoe, I. et al. CATH: increased structural coverage of functional space. Nucleic Acids Res 49, D266–D273 (2021).

23. Cox, J. & Mann, M. MaxQuant enables high peptide identification rates, individualized p.p.b.-range mass accuracies and proteome-wide protein quantification. Nat Biotechnol 26, 1367–72 (2008).

24. Choi, M. et al. MSstats: an R package for statistical analysis of quantitative mass spectrometry-based proteomic experiments. Bioinformatics 30, 2524–6 (2014).

25. Yu, G., Wang, L.G., Han, Y. & He, Q.Y. clusterProfiler: an R package for comparing biological themes among gene clusters. OMICS 16, 284–7 (2012).

26. Gadelha, H., Hernandez-Herrera, P., Montoya, F., Darszon, A. & Corkidi, G. Human sperm uses asymmetric and anisotropic flagellar controls to regulate swimming symmetry and cell steering. Sci Adv 6, eaba5168 (2020).

27. Perez-Riverol, Y. et al. The PRIDE database and related tools and resources in 2019: improving support for quantification data. Nucleic Acids Res 47, D442–D450 (2019).

