## Supplementary material for "*De novo* protein identification in mammalian sperm using high-resolution *in situ* cryo-electron tomography": Materials and Methods

All chemicals were purchased from Sigma-Aldrich unless otherwise noted.

### Sample preparation

Mouse sperm were collected from 10 to 16-week-old C57Bl/6J mice base on the published protocol[27]. Briefly, the sperm were stripped from vasa deferentia by applying pressure to cauda epididymis in 1 x Krebs buffer (1.2 mM $KH_2PO_4$, 120 mM NaCl, 1.2 mM $MgSO_4{\bullet}7H_2O$, 14 mM dextrose, 1.2 mM $CaCl_2{\bullet}2H_2O$, 5 mM KCl, 25 mM $NaHCO_3$). The sperm were washed and resuspended in ~100 µL Krebs buffer for the following experiments.

### Grid preparation

EM grids (Quantifoil R 2/2 Au 200 mesh) were glow discharged to be hydrophilic using an easiGlow system (Pelco). The grid was then loaded onto a Leica GP cryo plunger (pre-equilibrated to 95% relative humidity at 25 °C). The mouse sperm suspension was then mixed with 10-nm gold beads (Electron Microscopy Science, cat #25487) to achieve final concentrations at 2-6 x $10^6$ million cells/mL. EHNA (erythro-9-(2-hydroxy-3-nonyl)adenine) (Santa Cruz Biotechnology, CAS 51350-19-7) was added to a final concentration of 10 mM. Then 3.5 µL sperm mixture was added to each grid, followed by an incubation of 15 sec. The grids were then blotted for 4 sec and plunge-frozen in liquid ethane.

### Cryogenic focused ion beam (cryoFIB) milling

CryoFIB was performed using an Aquilos II cryo-FIB/SEM microscope (Thermo Fisher Scientific). A panorama SEM map of the whole grid was first taken at 377x magnification using an acceleration voltage of 5 kV with a beam current of 13 pA, and a dwell time of 1 µs. Targets with appropriate thickness for milling were picked on the map. A platinum layer (~10 nm) was sputter coated and a gas injection system (GIS) was used to deposit the precursor compound trimethyl(methylcyclopentadienyl) platinum (IV). The stage was tilted to 15-20°, corresponding to a milling angle of 8-13° relative to the plane of grids. FIB milling was performed using stepwise decreasing current as the lamellae became thinner

(1.0 nA to 30 pA, final thickness: ~300 nm). The grids were then stored in liquid nitrogen before data collection.

**Image acquisition and tomogram reconstruction**

Tilt series of mouse sperm were collected on a 300-kV Titan Krios transmission electron microscope (Thermo Fisher Scientific) equipped with a high brightness field emission gun (xFEG), a spherical aberration corrector, a Bioquantum energy filter (Gatan), and a K3 Summit detector (Gatan). The images were recorded at a nominal magnification of 26,000x in super-resolution counting mode using SerialEM[28]. After binning over 2 x 2 pixels, the calibrated pixel size was 2.612 Å on the specimen level. For each tilt series, images were acquired using a modified dose-symmetric scheme between -48° to 48° relative to the lamella with 4° steps and grouping of two images on either side (0°, 4°, 8°, -4°, -8°, 12°, 16°, -12°, -16°, 20°…)[29]. At each tilt angle, the image was recorded as movies divided into fourteen subframes. The total electron dose applied to a tilt series was 100 $e^-/Å^2$. The defocus target was set to be -2 to -5 μm.

All movie frames were corrected with a gain reference collected in the same EM session. Movement between frames was corrected using MotionCor2 without dose weighting[30]. Alignment of the tilt series and tomographic reconstructions were performed using Etomo[31]. The aligned tilt series were then CTF-corrected using TOMOCTF[32] and the tomograms were generated using TOMO3D[33] (bin4, pixel size: 10.448 Å).

In total, we started with 8 milling grids of mouse sperm and obtained 159 lamellae. Tilt series with no crystal ice were kept and further processed. In some cases, a significant part of the (9+2) axoneme was milled away and we only processed the ones with at least 5 doublets and also enough space including the central pair complex for subvolume averaging. In the end, the final reconstructions of the consensus averages were from 77 usable tomograms.

**Subvolume averaging**

Subsequent subvolume extraction, classification and refinement were all performed using RELION3[34] as reported previously. Briefly, subvolumes from the doublets were manually picked every 24 nm and extracted (pixel size: 15.672 Å, box size: 80 pixels, dimension: 125.376 nm). Initially, subvolumes were aligned to a map of non-treated mouse sperm doublet structure (EMDB-27444) lowpass filtered to 80 Å and the resulting map was used as the reference for further processing. Supervised 3D classification on radial spokes gave rise to four class averages of the 96-nm repeating units at four different registries. All four class averages were recentered at the base of Radial spoke 2 and re-extracted the same point (pixel size: 10.448 Å, box size: 120 pixels, dimension: 125.376 nm). All subvolumes were combined and aligned to one reference and duplicate subvolumes were removed based on minimum distance (< 40 nm). The remaining subvolumes were aligned to yield the consensus average for all nine doublets. These subvolumes were later remapped and sorted to calculate per-doublet averages. Subvolumes of the 96-nm repeating units were recentered on MIP features that repeat every 48 nm or 16 nm to obtain the coordinates of these subvolumes. The tomograms and coordinates were imported in RELION4 without binning (REF). Pseudo-subtomograms were extracted (pixel size: 2.612 Å, box size: 220 pixels, dimension: 57.464 nm) and the first round of Refine3D jobs yield the initial reference for the following refinement of geometric and optical parameters of the tilte series. TomoFrameAlign and CtfRefineTomo jobs were ran alternatively for two rounds and new pseudo-subtomograms were extracted. The same "Refine3D-TomoFrameAlign-CtfRefineTomo-TomoFrameAlign-CtfRefineTomo" process was repeated again and new pseudo-subtomograms were extracted. The final Refine3D job yield the reported maps. Further refinement did not improve the resolutions and quality of maps. In order to generate a map covering the entire 48 nm-repeating structure of the doublets, the run_data.star file from the final Refine3D job was shifted to opposite directions along the longitudinal axis by ~50 pixels and another round of refinement yield averages with the shifted registries. These three reconstructions were aligned to the reported 48 nm repeating structure of doublets from bovine trachea cilia and a composite map was generated to match the registry of the periodic structure.

To generate an average for the central pair complex, subvolumes were picked and extracted from the central pair complex every 16 nm (pixel size: 10.448 Å, box size: 100 pixels, dimension: 104.448 nm) (Extended Data Fig. 2b). Refinement of all subvolumes to an average of non-treated mouse sperm central apparatus (EMDB-27445) resulted in a central pair with only 16-nm repeating features. The alignment parameters were modified to reset all translations (x, y, z) to zero and then used for a second round of refinement with local search only. Focused classification was then performed on the microtubule-associated proteins on the C1 microtubule to separate the two populations of subvolumes (~50% each) corresponding to the central pair complex with 32-nm periodicity and an offset of 16 nm between them. The subvolumes were then re-centered and extracted on the same protein features for the two class averages. All subvolumes were aligned to the same reference using local searches. Duplicate subvolumes were removed based on minimum separating distance (< 30 nm). The remaining subvolumes were refined to generate the consensus average for the central pair complex.

The resolutions for maps were estimated based on FSC of two independently refined half datasets (FSC = 0.143). Local resolution maps for the consensus averages of doublets and the central pair complex of both human and mouse sperm were calculated by RELION4 and displayed in UCSF Chimera[35]. These local-resolution maps represent relative differences in resolution across the maps but the absolute values may not be exact. IMOD was used to visualize the tomographic slices[31]. UCSF Chimera was used to manually segment the maps for various structure features and these maps were colored individually to prepare the figures using UCSF ChimeraX[35-37].

**Unbiased matching of density maps to protein candidates**

MIP densities that are unique in sperm doublets were cropped from the corresponding maps. The PDB library of 21615 mouse proteins based on AlphaFold2 prediction was downloaded[13]. The unbiased matching was carried out using the COLORES program (Situs package)[38]. The matching was scored and ranked by the cross-correlation scores and top 200 hits were inspected individually with the target densities in UCSF Chimera. For the 4-helix bundle densities, CATH library, which curated non-redundant PDBs of published structural domains[22], was also used and no homologous proteins of SPACA9 was found.

### Model building

Model building was performed in Coot v0.9.8.1[39] and rigid body fitting was achieved using UCSF Chimera. Interpretation of the mouse sperm doublet map started with fitting of the atomic model of the bovine trachea doublet (PDB 7rro)[11]. Densities matching tubulins and 29 bovine MIPs in the bovine trachea doublet were found in the mouse sperm doublet map so all of these densities were considered to be formed by *Mus musculus* orthologs. These orthologs were identified using UniProt[40] or the NCBI protein database[41] based on the sequences of bovine proteins. The atomic models of the bovine MIPs were mutated to match the sequence of the mouse proteins using the Chainsaw plugin in Coot. The resulting models of individual proteins were then fit into the mouse sperm doublet map as rigid bodies. Model buildings for MIPs present in the mouse sperm doublets but absent in the bovine trachea (Tektin 5, CCDC105 and SPACA9) started with the AlphaFold2 predicted models and initial fitting was done by COLORES as described above. The loops that could not be traced were deleted. The orientations of helices are adjusted to fit into the densities in Coot.

### Sequence alignment and search for homologous proteins

Sequence alignment was performed using Clustal Omega[18] server and displayed in Jalview[42]. *Mus musculus* Tektin 1 sequence was used as input to search for Tektin homologs using the HHpred server[43].

### Biochemical extractions and mass spectrometry analyses of mouse sperm

For each biological replicate of the three, sperm from two mouse were washed by PBS and pelleted down at 2000 g for 5 min. Then E1-E5 buffers were used to extract proteins from the pellets [0.1 % Triton in PBS (E1), 0.6 M NaCl in PBS (E2), 0.6 M KCSN in PBS (E3), 8 M urea (E4) and 10% SDS (E5)]. For E1-E4, 100 µL of the buffer was added into the pellets the resuspension was mixed by pipetting up and down using a p200 pipette. Then the solution was incubated at room temperature for 10 min and the pellet was spun down at 21000 g for 10 min. For E5, after 10% SDS was added and mixed, the resuspension was heated at 95° for 5 min. After the pellets were spun down, the supernatant was taken as the extraction (E1-E5). 20 µL and 2 µL of the extractions were used for SDS-PAGE analyses, either stained with AcquaStain (Fisher Scientific, NCO170988) and blotted for an antibody against $\alpha$-tubulins (ThermoFisher Scientific, DM1A, #62204). The remaining extractions were used for mass spectrometry analyses.

**Mass spectrometry (MS)-based global protein abundance of mouse sperm**

Proteins in biochemical fractions E1, E2, E3, E4 and E5 from three biological replicates were reduced and alkylated in 4 mM final concentration tris (2-carboxyethyl) phosphine (TCEP) and 10 mM final concentration iodoacetamide by 20-minute incubation in the dark, after which excess iodoacetamide was quenched with 10 mM final concentration dithiothreitol (DTT). Proteins were then subjected to methanol chloroform precipitation. Briefly, 1 part sample was combined and vortexed sequentially with 4 parts methanol, 1 part chloroform, and 3 parts water for phase separation, after which samples were spun for 2 minutes at top speed (14,000 g) in a bench-top centrifuge (Centrifuge 5424R, Eppendorf). The upper phase was removed and discarded, and 4 parts methanol were combined and vortexed with the interphase and lower phase and subsequently centrifuged for 3 minutes at 14,000 g. The supernatant was removed and discarded, and the pellet was washed three times in 80% ice cold acetone followed by centrifugation for 3 minutes at 14,000 g. Extracted proteins were air dried, resuspended in 8 M urea buffer (8 M urea, 150 mM NaCl, 50 mM $NH_4HCO_3$, cOmplete Mini EDTA-free protease inhibitor (Roche, 11836170001)), and quantified using Bradford reagent (Sigma, B6916) following Coomassie (Bradford) Protein Assay Kit’s protocol (Thermo Fisher, 23200). Following quantification, protein samples were diluted 4-fold to 2 M urea concentration with 0.1 M $NH_4HCO_3$ pH 8, digested with trypsin (Promega, V5111) at a protease:protein ratio of 1:100 (weight/weight), and incubated overnight at 37ºC in a thermomixer at 750 rpm.

After tryptic digest, samples were acidified to pH <3 with 1% final concentration formic acid, and desalted for MS analysis using HPLC-grade reagents and 100 µL OMIX C18 tips (Agilent Technologies, A57003100) according to the manufacturer’s protocol with the following adjustments. Briefly, OMIX tips were conditioned by sequential washes of 100% acetonitrile and 50% acetonitrile, 0.1% formic acid, and equilibrated with two washes of 0.1% formic acid. Peptides were bound to the C18 polymer by repeated pipetting, subsequently washed three times with 0.1% formic acid, and sequentially eluted in 50% acetonitrile, 0.1% formic acid followed by 90% acetonitrile, 0.1% formic acid. Peptides were dried by vacuum centrifugation (CentriVap Cold Trap, Labconco) and stored at -80ºC until MS analysis.

Digested, desalted peptides were resuspended to 0.125-2 µg/µL final concentration in 2% acetonitrile, 0.1% formic acid. 1-2 µL were injected in technical singlet onto an Easy-nLC 1200 (Thermo Fisher Scientific) interfaced via a nanoelectrospray source (Nanospray Flex) coupled to an Orbitrap Fusion Lumos Tribrid mass spectrometer (Thermo Fisher Scientific). Peptides were separated on a PepSep reverse-phase C18 column (1.9 µm particles, 1.5 µm x 15 cm, 150 µm ID) (Bruker) with a gradient of 5-88% buffer B (0.1% formic acid in acetonitrile) over buffer A (0.1% formic acid in water) over a 100-minute data acquisition. Spectra were acquired continuously in a data-dependent manner. One full scan in the Orbitrap (scan range 350-1350 m/z at 120,000 resolution in profile mode with a custom AGC target and maximum injection time of 50 milliseconds) was followed by as many MS/MS scans as could be acquired on the most abundant ions in 2 seconds in the dual linear ion trap (rapid scan type with fixed HCD collision energy of 32%, custom AGC target, maximum injection time of 50 milliseconds, and isolation window of 0.7 m/z). Singly and unassigned charge states were rejected. Dynamic exclusion was enabled with a repeat count of 1, an exclusion duration of 25 seconds, and an exclusion mass width of ±10 ppm. Liquid chromatography (LC) and MS acquisition parameters are reported in (Supplementary Table S1).

Raw MS files were searched using MaxQuant (version 1.6.3.3) against a database of the mouse proteome (SwissProt Mus musculus reviewed protein sequences, downloaded 07 May 2022) with a manual addition to include mouse piercer of microtubule wall 2 protein (protein sequence from NCBI Reference Sequence NP_001185718.1, manually assigned the UniProt identifier "ZCC15orf65" in our database after its bovine homolog)[23]. MaxQuant settings were left at default, with the following exceptions: LFQ was enabled with skip normalization enabled; and match between runs was enabled with a 1.5-minute matching time window and 20-minute alignment window. Trypsin (KR|P) was selected and allowed up to two missed cleavages, and variable and fixed modifications were assigned for protein acetylation (N-terminal), methionine oxidation and carbamidomethylation.

Statistical analysis of protein quantitation was completed with R Bioconductor package artMS (version 1.14.0) (doi: 10.18129/B9.bioc.artMS) and its function artmsQuantification, which is a wrapper around the R Bioconductor package Mass Spectrometry Statistics and Quantification (MSstats) (version 4.4.0) as follows[24] (Supplementary Table S2). Peptide intensities from the MaxQuant evidence file were summarized to protein intensities using the MSstats function dataProcess with default settings. The differences in log2-transformed intensity between biochemical fractions were scored using the MSstats function groupComparison, which fits a single linear model for each protein with a single categorical variable for condition, or fraction in our case. From these models, MSstats reports pairwise differences in means between conditions as log2 fold change (log2FC) with a p-value based on a t-test assuming equal variance across all conditions, and reports adjusted p-values using the false discovery rate (FDR) estimated by the Benjamini-Hochberg procedure. Proteins with significant changes in abundance between fractions were defined as: (1) absolute(log2FC) > 1; and (2) adjusted p-value < 0.05. Proteins with significant changes in abundance were tested for enrichment of Gene Ontology terms (Supplementary Table S3). The over-representation analysis was performed using the enricher function from R package clusterProfiler (version 4.4.1)[25]. Gene Ontology (GO Biological Process, Molecular Function and Cellular Component) terms and annotations were obtained from the R annotation package org.Mm.eg.db (version 3.15.0). From among all significantly enriched terms, we selected a set of non-redundant terms following a clustering procedure. We first constructed a term tree based on distances (1-Jaccard Similarity Coefficients of shared genes in KEGG or GO) between the significant terms. The term tree was cut at a specific level (h = 0.99) to identify clusters of non-redundant gene sets (Supplementary Table S4). For results with multiple significant terms belonging to the same cluster, we selected the most significant (lowest adjusted p-value) term.
